# Enhanced proconvulsant sensitivity, not spontaneous rapid swimming activity, is a robust correlate of *scn1lab* loss-of-function in stable mutant and F0 crispant hypopigmented zebrafish expressing GCaMP6s

**DOI:** 10.1101/2025.01.15.633275

**Authors:** Christopher Michael McGraw, Cristina M. Baker, Annapurna Poduri

## Abstract

Zebrafish models of genetic epilepsy benefit from the ability to assess disease-relevant knock-out alleles with numerous tools, including genetically encoded calcium indicators (GECIs) and hypopigmentation alleles to improve visualization. However, there may be unintended effects of these manipulations on the phenotypes under investigation. There is also debate regarding the use of stable loss-of-function (LoF) alleles in zebrafish, due to genetic compensation (GC). In the present study, we applied a method for combined movement and calcium fluorescence profiling to the study of a zebrafish model of *SCN1A*, the main gene associated with Dravet syndrome, which encodes the voltage-gated sodium channel alpha1 subunit (Nav1.1). We evaluated for spontaneous and proconvulsant-induced seizure-like activity associated with *scn1lab* LoF mutations in larval zebrafish expressing a neuronally-driven GECI (elavl3:GCaMP6s) and a *nacre* mutation causing a common pigmentation defect. In parallel studies of stable *scn1lab^s^*^552^ mutant*s* and F0 crispant larvae generated using a CRISPR/Cas9 multi-sgRNA approach, we find that neither stable nor acute F0 larvae recapitulate the previously reported seizure-like rapid swimming phenotype nor does either group show spontaneous calcium events meeting criteria for seizure-like activity based on a logistic classifier trained on movement and fluorescence features of proconvulsant-induced seizures. This constitutes two independent lines of evidence for a suppressive effect against the *scn1lab* phenotype, possibly due to the GCaMP6s-derived genetic background (AB) or *nacre* hypopigmentation. In response to the proconvulsant pentylenetetrazole (PTZ), we see evidence of a separate suppressive effect affecting all conspecific larvae derived from the stable *scn1lab^s^*^552^ line, independent of genotype, possibly related to a maternal effect of *scn1lab* LoF in mutant parents or the residual TL background. Nonetheless, both stable and F0 crispant fish show enhanced sensitivity to PTZ relative to conspecific larvae, suggesting that proconvulsant sensitivity provides a more robust readout of *scn1lab* LoF under our experimental conditions. Our study underscores the unexpected challenges associated with the combination of common zebrafish tools with disease alleles in the phenotyping of zebrafish models of genetic epilepsy. Our work further highlights the advantages of using F0 crispants and the evaluation of proconvulsant sensitivity as complementary approaches that faithfully reflect the shared gene-specific pathophysiology underlying spontaneous seizures in stable mutant lines. Future work to understand the molecular mechanisms by which *scn1lab*-related seizures and PTZ-related hyperexcitability are suppressed under these conditions may shed light on factors contributing to variability in preclinical models of epilepsy more generally and may identify genetic modifiers relevant to Dravet syndrome.

## 1. Introduction

Zebrafish have emerged as a powerful model of chemical and genetic seizures^1,2^ with many advantages, including large clutch size, rapid external development, and high genetic conservation with higher vertebrates for modeling disease. Zebrafish are amenable to genetic manipulation by CRISPR/Cas9 gene editing, enabling the study of gene-specific loss of function through the generation of stable gene knockout lines as well as acute crispant knockouts in the F0 generation. In addition, a number of tools enable the advanced study of zebrafish brain activity, such as genetically encoded calcium indicators (GECIs; for example, GCaMP6s^3^) and pigmentation mutants (such as *nacre*^4^) that improve visualization for live imaging. The small size of larval zebrafish also makes them suitable for higher throughput screening^2,5^. Recently we described a platform^6^ that combines movement and fluorescence data from unrestrained zebrafish for the evaluation of chemically induced seizures using 96-well format, but its application to genetic models of epilepsy has not yet been reported.

Despite its strengths, the study of disease in zebrafish is challenged by at least two factors. First, the genetic backgrounds of laboratory zebrafish (AB, TL, and others^7^) are often not carefully reported or controlled across experiments by zebrafish researchers^8^. Genetic modifiers, which refer to discrete genetic factors capable of “modifying” the severity or penetrance of a phenotype, have been well-described in association with different inbred mouse strains (for example, in epilepsy models^9^), and recognized to alter phenotypes between wild-type zebrafish lines^7,8^, but their significance in zebrafish models of disease or epilepsy remains considerably less well-explored. This is a critical issue for the rigor and reproducibility of zebrafish studies^8^ because if genetic tools generated on one zebrafish background harbor modifiers that affect the phenotype of mutant alleles generated on a different zebrafish background, the fundamental logic by which any mechanistic claim derived from the use of these tools may be compromised.

Second, whether any stable loss-of-function allele will demonstrate a phenotype in larval zebrafish is somewhat unpredictable, owing to the effects of genetic compensation (GC), about which there have been increasing reports in zebrafish^10^. GC refers to the many processes by which the effects of a genetic perturbation are functionally balanced by compensatory changes in the regulation of other genes, the details of which remain incompletely understood. One type of GC termed transcriptional adaptation (TA) has been shown to be triggered by premature termination codons (PTCs) through the nonsense mediated decay (NMD) pathway, and lead to upregulation of genes with sequence homology including paralogs^11^. The TA response appears to be epigenetically passed on to genetically wild-type offspring^12^, though other modes of GC can also exert influence on progeny independent of larval genotype if they perturb the levels of mRNA/proteins present in the maternal gametes via so-called maternal effects^13^. As an alternative to stable lines, gene-specific acute F0 crispants -- generated by microinjection of Cas9 ribonucleoproteins and guide RNA (gRNA) into fertilized embryos – are often used and in some instances appear to have stronger phenotypes than stable alleles^14^, perhaps in part by avoiding maternal effects.

Here we apply the combined movement and fluorescence approach to the characterization of genetic epilepsy models through the example of the epilepsy gene *SCN1A.* The gene *SCN1A* encodes the voltage-gated sodium channel alpha1 subunit (Na_v_1.1), and human variants in *SCN1A* are associated with Dravet syndrome (DS), a severe developmental epileptic encephalopathy characterized by drug-refractory seizures^15^. Zebrafish with disruptions in the homologous gene *scn1lab* have been studied as models of DS^5,16–18^ and recapitulate key features of the disease, including seizures and their response to anti-seizure medication. Specifically, the homozygote larvae from the well-described *scn1lab^s^*^552^ (Didy) allele (harboring a Met-to-Arg missense mutation in exon 18^16,19^) have demonstrated seizure-like activity across multiple modalities including assays of locomotor behavior (bursts of rapid swimming, >20mm/sec), tectal recordings of local field potential (high amplitude frequent epileptiform discharges), as well as calcium fluorescence (seizure-like bursts^20^). For these reasons, the phenotype associated with *scn1lab* fish is considered a gold-standard control for evaluating methods of seizure detection.

In the present study, our goal was to benchmark the combined movement and fluorescence profiling approach^6^ for identifying seizure-like activity in models of genetic epilepsy. Towards this end, we evaluated spontaneous and proconvulsant-induced seizure-like activity associated with *scn1lab* loss-of-function (LoF) mutations in larval zebrafish expressing a neuronally-driven, genetically encoded calcium indicator (elavl3:GCaMP6s^3^) in combination with a common hypopigmentation defect (*nacre*^4^). We chose the stable *scn1lab^s^*^552^ allele as a positive control. To assess the phenotypic similarity between *scn1lab^s^*^552^ and acute CRISPR/Cas9 mediated knock-out, we also generated acute *scn1lab* F0 crispant larvae using a multi-sgRNA approach. We expected to recapitulate the well-documented seizure-related phenotypes associated with *scn1lab* LoF, but instead we observe that neither stable nor acute F0 fish recapitulate the previously reported spontaneous rapid swimming phenotype, suggesting a suppressive effect under these conditions possibly related to genetic background or other factors. Similarly, neither group showed any evidence for spontaneous calcium events meeting criteria for seizure-like activity based on a machine learning classification trained on movement and fluorescence features of proconvulsant-induced seizures^6^. In addition, we observe *totally opposite* effects of *scn1lab* LoF on several event-related parameters between the two approaches, with stable *scn1lab* mutant larvae showing elevations in maximum velocity, average distance, and calcium event rate versus crispant F0s showing reductions relative to their respective conspecific controls. We also see markedly reduced PTZ sensitivity in the *scn1lab^s^*^552^ line, affecting mutant and wild-type conspecifics, which may be due to genetic background or a maternal effect of parental *scn1lab* LoF. Despite these limitations, both stable and F0 crispant fish show enhanced sensitivity to PTZ relative to conspecifics, suggesting that proconvulsant sensitivity may be a more robust readout of *scn1lab* LoF and perhaps other epilepsy-related genes under these conditions.

## 2. Materials and methods

### 2.1. Zebrafish maintenance

GCaMP6s^22^ zebrafish (*Danio rerio*) with *nacre* pigmentation deficit used for all experiments were obtained on AB background as TG(*elav3*::Gcamp6s); *mitfa*^w2/w2^ (abbreviated, *GCaMP6s*; generous gift from Florian Engert, Harvard University). Zebrafish were maintained by in-crosses, and larvae periodically selected for “high” GCaMP6s fluorescence and nacre phenotype. The *scn1lab^s^*^552^ (Didy^16^) line was obtained as a generous gift from H. Baier (Max Plank Institute of Neurobiology) on TL background. Heterozygote *scn1lab^s^*^55a/+^ fish were crossed to *Gcamp6s*, and adult F1 fish were in-crossed to obtain F2 *Gcamp6s*; nacre for experiments, and are anticipated to be roughly 50:50 AB:TL. All fish were maintained on a 14H:10H day-night cycle. All procedures were approved by BCH Animal Welfare Assurance (IACUC protocol #00001775).

### 2.2. Genotyping *scn1lab*^s552^ line

For *scn1lab^s^*^552^, the following primers were used for PCR to generate a 298bp fragment: *scn1lab*-diddy-FW, GCTGTGTGATGAGGTTTCAGT; *scn1lab*-diddy-RV, CTGTTAGACAGAAATTGGGGG. SnapGene was used to inspect the chromatogram from Sanger sequencing and to identify larvae with the c.T>G mutation at the sequence TTCA<T/G>GATT (Supplementary Fig 1) in Exon 18 (Ensembl ID, ENSDARE00000666203).

### 2.3. Crispant *scn1lab* generation

#### 2.3.1. sgRNA design and synthesis

Three sgRNAs were designed using the online CHOPCHOP tool (V2^21^) with default settings targeting exonic regions of the zebrafish gene *scn1lab,* selecting only sgRNA with no predicted off-target activity (MM0-MM3 = 0), and efficiency >0.6. The selected sgRNA corresponded to ranks 1, 3, 7, and had predicted efficiency scores of 0.74, 0.73, and 0.71, respectively.

*scn1lab* sgRNA1 GGTTACAGTACCGATAGCGG exon 16

*scn1lab* sgRNA2 GTTTAGAGCCGGCCAAGAAG exon 16

*scn1lab* sgRNA3 TATTCGCCCCCCTGGAGAGG exon 17

Synthetic sgRNA with chemical modifications 2’-O-Methyl at 3 first and last bases and 3’ phosphorothioate bonds between first 3 and last 2 bases were ordered from Synthego (Redwood City, CA)

#### 2.3.2. Microinjection

Upon receipt, sgRNA were diluted to 1000ng/uL and mixed 1:1:1 before freezing at -80degC. For microinjection, pooled sgRNA were thawed on ice, and mixed with sterile H20 and phenol red to maintain a final gRNA concentration of 250ng/uL. Embryos were derived from timed in-cross matings from GCaMP6s;*nacre* parents. Microinjections (2nL, or 150 micron diameter) into yolk sac of fertilized embryos at the one-cell stage were performed with all injections completed within 20-40 minutes of fertilization.

#### 2.3.3. Assessment of CRISPR efficiency

To control for subject-specific differences in cutting efficiency, we assayed cutting at *scn1lab* loci by PCR amplification, followed by Sanger sequencing and ICE analysis^22^. Genomic DNA from larvae (dpf 5-6) was obtained by sodium hydroxide digestion (NaOH 50mM final concentration), heated incubation at 90degC x 1-2hrs, followed by neutralization with 1/10^th^ volume 1M Tris-HCl (pH 8). PCR was performed to generate amplicons with the following primer pairs corresponding to each sgRNA guide sequence. The F/R primers for each reaction are as follows: 1) “*scn1lab* sgRNA1 PCR F” AAGGACTATCTGAAGGAGGGCT; “*scn1lab* sgRNA1 PCR R” TCTCTCCGACACTGAAACAAGA (product size 288bp); 2) “*scn1lab* sgRNA2 PCR F” ACAGAAAGGTATCGCTCTGGTC; “*scn1lab* sgRNA2 PCR R” ACATGTAGTCGCCTTCCTCAAT (product size 260bp); 3) “*scn1lab* sgRNA3 PCR F” ACCTGTCGATACGGTTCTCAGT; “*scn1lab* sgRNA3 PCR R” CACTAAATTGGCCAGTGTTTCA (product size 268bp).

Each amplicon was Sanger sequenced (GeneWiz) with F or R primer, and the percentage of cutting associated with inferred knock-out (“KO score”) obtained using the Synthego Inference of Crisper Edits (ICE) tool^22^. For simplicity, injected larvae were stratified into 3 categories, NO CUT (0%) vs LOW (0-50%) vs HIGH (>50%) based on the KO score from the first sgRNA reaction, and compared to uninjected conspecific controls.

### 2.4. PTZ concentration escalation

PTZ (Sigma; stored -20degC) was prepared fresh in sterile fish water (Instant Ocean) to a stock concentration 26mM, then diluted to intermediate concentrations. For serial concentration escalation experiments, a standard 10uL volume from PTZ Stock 1 (25.7mM) was added to each well of a 96-well plate (100uL starting volume per well) to yield 2.5mM, followed by an additional 10uL from PTZ Stock 2 (152.5mM; 110uL starting volume per well) to yield 15mM final concentration (final well volume, 120uL). For experiments involving anti-seizure drug pretreatment, anti-seizure drugs were administered in a standard 10uL volume. Following baseline recording, a standard 10uL volume from PTZ Stock 1 (30mM) was added to each well of a 96-well plate (110uL starting volume per well) to yield 2.5mM, followed by an additional 10uL from PTZ Stock 2 (165mM; 120uL starting volume per well) to yield 15mM final concentration (final well volume, 130uL). Pipetting was performed manually with a multi-channel pipettor. Three sequential 30-minute recordings were performed during baseline, PTZ 2.5mM, and PTZ 15mM conditions, respectively.

### 2.5. Calcium fluorescence imaging

Imaging was performed as previously reported^6^. Briefly, individual unrestrained larval zebrafish (dpf 5) are placed into wells of an optical 96-well plate (Greiner 655076) in 100uL sterile fish water (Instant Ocean) and imaged using the FDSS7000EX fluorescent plate reader (Hamamatsu; software version 2). Specimens are illuminated by a Xenon light source passed through a 480nm filter. Epifluorescence from below the specimen is filtered (540nm) and collected by EM-CCD, allowing all wells to be recorded simultaneously. Data is collected as 256×256, 16-bit image at ∼12.6 Hz (79 msec interval), 2×2 binning, sensitivity setting = 1. Image data was extracted from the .FLI file using ImageJ or MATLAB based on the following parameters: 16-bit unsigned, 256×256, offset 66809, gap 32 bytes. Analysis was performed in MATLAB to extract position, linear and angular velocity, and changes in calcium fluorescence using a moving average deltaF/F0 method.

### 2.6. Analysis of calcium fluorescence data

Analysis was performed as previously reported^6^. Briefly, an algorithm to track changes in calcium activity using a “moving delta F/F0” was devised in MATLAB. The initial 256×256 time-series is segmented into individual wells (∼14 x 14 pixels, ∼0.513mm per pixel) based on a pre-specified plate map. For each well, the n x m x t time-series is expanded to 2n x 2m x t using bicubic interpolation before further processing. Calcium transients are detected based on the normalized instantaneous average fluorescence for the area of the fish body within the well by the following formula: (average Ffish(t) -F0)/F0, where F0 = average Ffish (averaged over each pixel, for each time sample), and smoothed with a 1000-sample (∼79 seconds) boxcar moving average. Fish x,y position is tracked based on a weighted centroid, and linear and angular velocity estimated. The minimum detectable change in the position of a larval zebrafish is estimated to be 0.256mm, corresponding to the size of one pixel after interpolation. For detecting significant fluctuations in calcium fluorescence (referred to as calcium events), the F/F0 time-series is further smoothed with a 25-sample (∼1.975 seconds) boxcar moving average. Calcium events are initially detected from the smoothed delta F/F0 time-series by identifying peaks that exceed an empirically determined permissive threshold (0.05), while the start and end of each event is identified by the zero-crossing of the smoothed 1st derivative. Subsequently, multiple per-event measurements are obtained for each event based on combined movement and fluorescence measurements, including: (1) MaxIntensity_F_centroid: the maximum fluorescence value of the detected fish during an event; (2) MaxIntensity_F_F0_centroid: the maximum delta F/F0 value within the boundary of the detected fish during an event; (3) distance_xy_mm: total distance moved during an event in millimeters; (4) duration_sec: elapsed time in seconds; and (5) total_revolutions: number of complete circles traveled by the fish during an event.

In addition, multiple per-fish measurements are obtained, including: (1) maxRange_2: the maximum fluorescence value observed during the recording; (2) totalCentroidSize_mode_mm2: the total area in square millimeters of the detected fish that exceeded a hard-coded threshold above sensor noise, which relates to the brightness of the fish.

### 2.7. Supervised machine learning for event classification

To differentiate calcium events related to seizure-like activity from other causes of calcium fluctuation, we used a previously described logistic classifier having been fit to a combination of event-level and fish-level features in R using elastic net regression via the *train*() function (R package, *caret*) and the *<u>glmnet</u>*method (R package, *glmnet*) as previously published^6^ and publicly available (http://doi.org/10.17605/OSF.IO/TNVUJ). This model (referred to as the “PTZ M+F” model) was previously trained on calcium events from PTZ-induced seizures (15mM) versus baseline conditions, and distinguishes seizure-like activity from non-seizure-like activity with high accuracy^6^. The model was used to classify calcium events from *scn1lab* animals as seizure-like or non-seizure-like using R. Fish lacking minimum fluorescence criteria (mode of fluorescence area < 0.05 mm2) were excluded from analysis.

### 2.8. Bootstrap simulation to identify optimal replicate number

Bootstrap simulations were conducted in R using custom code and the *rep_sample_n()* function (R package, *moderndive*) as described in the main text. The robust strictly standardized mean difference (RSSMD) between target(1) and background(2) is calculated^23^ as: 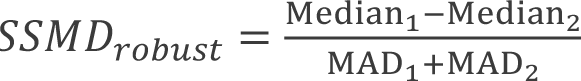 the median absolute deviation (MAD) is defined as: *MAD* = Median(| X − Median(X)|).

### 2.9. Statistical analysis

Unless otherwise indicated, the statistical significance of group-wise differences was assessed using the non-parametric Wilcoxon rank sum test in Prism (v10.0.3, GraphPad). The false discovery rate (FDR) for multiple comparisons was controlled using the two-stage linear step-up procedure of Benjamini, Krieger, and Yekutieli to maintain family-wise alpha = 0.05. Only adjusted p-values after FDR correction are reported.

## 3. Results

### 3.1. Loss-of-function mutations in the scn1lab gene are not associated with spontaneous seizure-like activity and have opposite effects on velocity, distance traveled, and event rate in scn1lab F0 versus scn1lab^s552^ hypopigmented transgenic GCaMP6s zebrafish

We began by looking at the pattern of movement and calcium fluorescence changes in freely moving *scn1lab* fish at baseline (**Fig 1A**) using a specialized fluorescent plate-reader. Stable *scn1lab^s^*^552^ larvae were generated from an incross of mutant parents to yield conspecific controls (hereafter referred to as WT, HET, or HOM). Acute F0 crispant larvae were generated by CRISPR/Cas9, combining 3 sgRNA targeting exons 16-17 of *scn1lab* (**Fig 1B**), and stratified by cutting efficiency (NO CUT (0%) vs LOW (0-50%) vs HIGH (>50%)) versus uninjected conspecific controls. To lend insight into the presence of clutch-specific effects, we also compared these results to two separate age-matched cohorts of wild-type fish derived from incross of GCaMP6s;*nacre* fish (referred to as “WT1” and “WT2”). All tested larvae had *nacre* hypopigmentation phenotype and neuronally expressed GCaMP6s.

**Figure 1.**
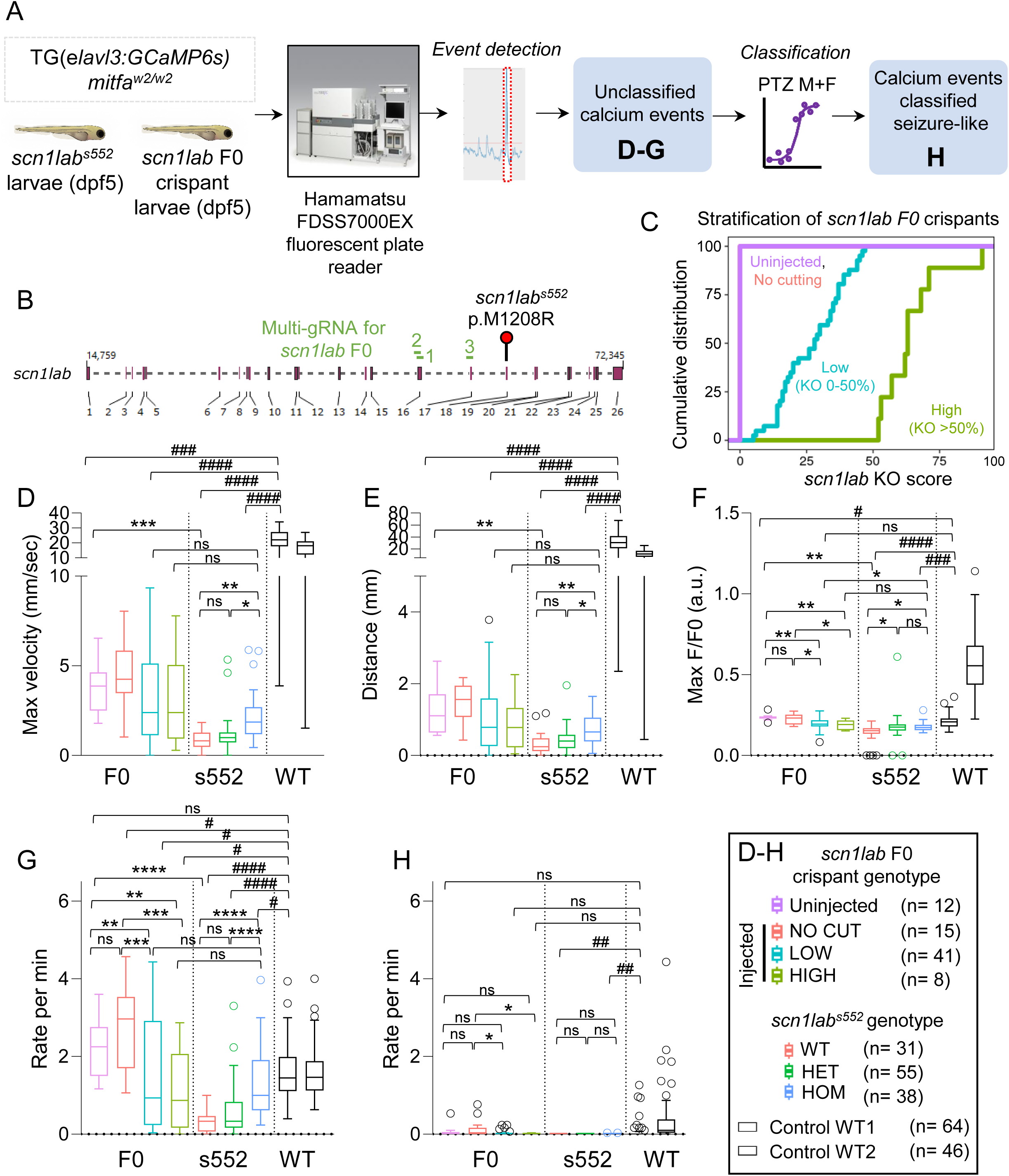
Loss-of-function mutations in the *scn1lab* gene are not associated with spontaneous seizure-like activity and have opposite effects on velocity, distance traveled, and bout rate in *scn1lab* F0 versus *scn1lab^s^*^552^ hypopigmented transgenic GCaMP6s zebrafish. (**A**) Schematic overview of experiment to observe spontaneous activity of unrestrained larvae, without classification and following classification with a logistic classifier trained to detect seizure-like activity using both movement and fluorescence related features. (**B**) Organization of the *scn1lab* locus in zebrafish, with numbered exons. The position of gRNA sequences used to generate *scn1lab* F0 crispants is shown in green. The position of the M1208R mutation in exon 18 of the *scn1lab^s^*^552^ allele is shown as a red lollipop. (**C**) Cumulative distributions of cutting at *scn1lab* by ICE KO score, derived from Sanger sequencing. (**D-F**) Parameters from spontaneous activity of unrestrained larvae, derived from combined movement and fluorescence profiling, including maximum velocity (D), average distance (E), and maximum normalized calcium fluorescence (dF/F0; F). (**G-H**) Rate of calcium events observed without classification or filtering (G) and after applying the PTZ M+F classifier (H). Data are group-wise Tukey box-plots of subject-level averages of all subject-specific events. Animal numbers for each group are indicated in the legend. Reported p-values are derived from Wilcoxon rank sum test, and adjusted by false-discovery rate (FDR) correction. The hash (#) symbol is used to indicate comparisons with a WT control cohort (GCaMP6s; *nacre*) not derived from the experimental cross. * or #, adj P<0.05. ** or ##, adj P<0.01. *** or ###, adj P<0.001. **** or ####, adj P<0.0001.

First, we observed marked suppression of average maximum velocity in both lines (**Fig 1D**). Wild-type controls and mutants from acute F0 and stable *s552* cohorts had significantly reduced velocity (max velocity, uninjected: median 3.87 mm/sec, IQR 2.5-4.6; WT: Median 0.81 mm/sec, IQR 0.47-1.28) relative to heterospecific wild-type controls (control WT1: median 22.07 mm/sec, IQR 17.83-27.39’; adj. P = 0.0002 (vs uninjected), adj. P <0.0001 (vs. WT), Wilcoxon rank sum with false discovery rate (FDR) correction). Prior studies regarding the *scn1lab^s^*^552^ line^5,16^ have reported that the presence of high-velocity movements (greater than or equal to 20 mm/s) are specific for paroxysmal whole body convulsions (referred to as stage III seizures) and that the activity is highly penetrant, and limited to *scn1lab^s^*^552^ HOM larvae, never being observed in conspecific controls. In contrast, our data suggests that high-velocity locomotor activity does occur in unaffected animals and is not highly prevalent in *scn1lab* mutants, at least not on this background or under our experimental conditions.

Second, in this context, we observed no rapid swimming phenotypes in LOW/HIGH F0 crispants or in stable *s552* HOM fish (**Fig 1D**), either by absolute criteria (>20mm/sec) or relative to conspecific controls. In fact, there appear to be opposite effects of *scn1lab* LoF in each of these scenarios, with *s552* HOM fish showing increased maximum velocity (median 1.87 mm/s, IQR 1.19-2.70), relative to conspecific controls (WT: 0.81 mm/s, IQR 0.48-1.30, adj P = 0.0049; HET: 0.99 mm/s, IQR 0.73-1.3, adj P = 0.014). Meanwhile, LOW/HIGH cutting F0 crispants instead show modest reductions in velocity (LOW: Median 2.39 mm/s, IQR (1.13-5.13), adj P = 0.12 (versus on injected), 0.07 (versus NO CUT); HIGH: 2.39 mm/s, IQR 0.94-5.05, adj P = 0.19 (versus injected), 0.15 (versus NO CUT)) versus conspecific controls. This observation is further corroborated by the fact that heterospecific controls are significantly different between these cohorts (uninjected versus *s552* WT, adj P = 0.005) while F0 crispant and stable *s552* HOM are not significantly different (LOW versus HOM: adj P = 0.228; HIGH versus HOM: adj P = 0.297). These findings are similar to those observed for average distance per bout (**Fig 1E**). The same effects are also seen in normalized calcium fluorescence (**Fig 1F**), but the differences between cohorts and that of heterospecific WT controls is less dramatic in this context.

In our previous work, we have demonstrated how the rate of calcium events is a quantitative measure of seizure-like activity^6^, therefore we also assessed the rate of calcium events in *scn1lab* fish (**Fig 1G**). Here again, and to an even stronger degree, we observed that the effect of *scn1lab* loss of function on the rate of unclassified calcium events is opposite between the lines. In *s552* HOM fish, the rate of events is elevated (median 1/min, IQR 0.62-1.9) versus conspecifics (WT: 0.33/min, IQR 0.067-0.467, adj P <0.0001; HET: 0.33/min, IQR 0.167-0.833, adj P <0.0001). Meanwhile, LOW and HIGH cutting F0 crispants instead showed reductions (LOW: 0.933/min, IQR 0.23-2.92; HIGH: 0.867/min, IQR 0.167-2.07) versus conspecifics (uninjected: 2.25/min, IQR 1.5-2.76, adj P = 0.0025 (versus LOW), 0.0034 (versus HIGH); NO CUT: 2.97/min, IQR 1.7-3.53, adj P = 0.001 (versus LOW), 0.006 (versus HIGH)). Nevertheless, LOW/HIGH F0 crispant and *s552* HOM larvae are actually quite similar (LOW versus HOM, adj P = 0.33; HIGH versus HOM, adj P = 0.16), whereas heterospecific controls are markedly different (uninjected versus WT, adj P < 0.0001; NO CUT versus HET, adj P < 0.0001).

Using a previously described logistic classifier^6^ (referred to as “PTZ M+F” classifier) trained on movement- and fluorescence-related features of seizure-like events induced by the proconvulsant GABA_A_R antagonist pentylenetetrazole (PTZ), we also assessed the rate of calcium events classified as seizure-like in *scn1lab* larvae (**Fig 1H**). Virtually no events are classified as seizure-like in either F0 crispant or stable *s552* larvae, suggesting that if seizure-like activity is occurring in *scn1lab* animals, it is distinct from that associated with PTZ.

To address this possibility, we trained a classifier using elastic net logistic regression on events from *s552*-HOM fish versus WT to determine whether the spontaneous unclassified calcium events in HOM animals might be comprised of events with milder “seizure-like” features (**Supplemental Methods; Supplemental Figure 2**). This classifier did not achieve high accuracy (**SFig 2B**; AUC-ROC 0.75; AUC-PRG, 0.17; F1 score, 0.696), and although it did confirm two classes of events (type 0 vs type 1) enriched in HOM versus WT animals, respectively (**SFig 2D,F**) – the former associated with small but significant elevations in the max velocity (**SFig 2I**) and distance per event (**SFig 2G**) relative to conspecifics – the type 0 events were not sufficiently different from events observed under physiological conditions in heterospecific wild-type control animals to justify calling them “seizure-like” with confidence. In addition, applying the *scn1lab s552* HOM classifier to F0 crispant animals yielded anomalous results (**SFig 2K**), with NO CUT controls showing elevated rates of type 0 events relative to LOW and HIGH groups, suggesting again that type 0 events are a variant of normal physiological events. Alternately, the features accessible by combined movement and fluorescence profiling may be insufficient to distinguish genetic seizures accurately, at least in the setting of the unexpected suppressive phenomenon that we observed here.

In summary, although the mechanism for these findings remains unclear, it is surprising not to see conservation of the previously reported rapid swimming phenotype either in stable *s552* line or in the acute F0 line. The observation that totally opposite gene-specific phenotypes are possible in response to *scn1lab* LoF – a well-characterized epilepsy gene – is also surprising and problematic, for efforts to compare phenotypes between stable and F0 crispant lines. At a minimum, we conclude that these parameters alone may be unreliable for the characterization of spontaneous seizure-like activity in the context of combined movement and fluorescence profiling in novel mutant larvae under these conditions.

### 3.2. Loss-of-function mutations in the scn1lab gene are associated with enhanced susceptibility to GABA_A_R antagonist pentylenetetrazole (PTZ) in both scn1lab F0 and scn1lab^s552^ hypopigmented transgenic GCaMP6s zebrafish

We asked whether *scn1lab* LoF affects proconvulsant sensitivity using the combined movement and fluorescence profiling approach (**Fig 2A**) and a serial concentration escalation paradigm using low-concentration PTZ (2.5mM), followed by high-concentration PTZ (15mM), as previously reported^6^.

**Figure 2.**
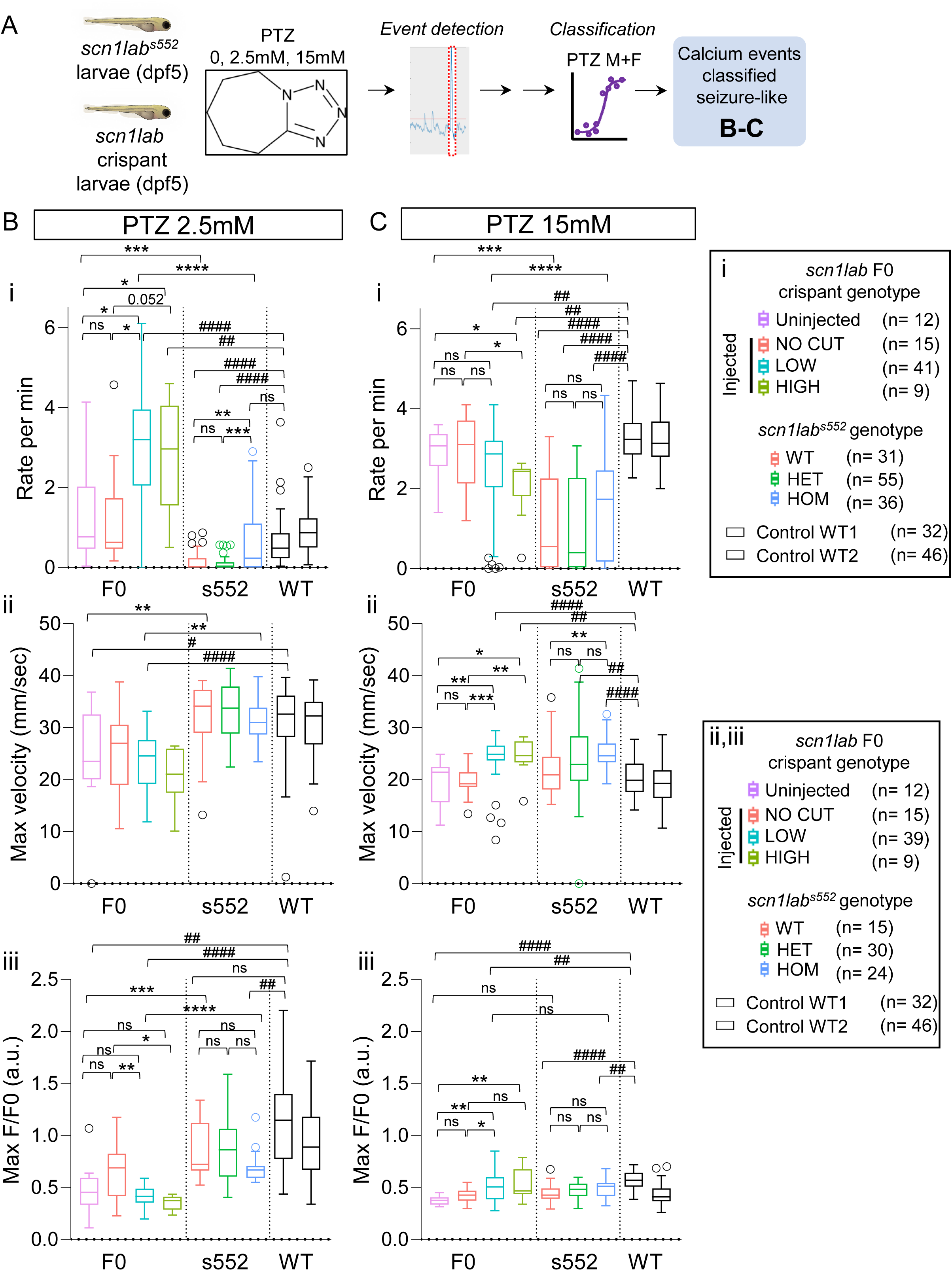
Loss-of-function mutations in the *scn1lab* gene are associated with enhanced susceptibility to GABA_A_R antagonist, pentylenetetrazole (PTZ) in both *scn1lab* F0 and *scn1lab^s^*^552^ hypopigmented transgenic GCaMP6s zebrafish. (A) Schematic overview of PTZ-concentration escalation experiment in unrestrained larvae, followed by classification of events using the PTZ M+F classifier. (B-C) Parameters derived from proconvulsant-induced activity of unrestrained larvae at low-concentration (2.5mM; B) and high-concentration (15mM; C) PTZ. Data are group-wise Tukey box-plots of subject-level averages of all subject-specific events. Animal numbers for each group are indicated in the legend according to subpanel; only animals with detectable events were analyzed in subpanels ii-iii. Reported p-values are derived from Wilcoxon rank sum test, and adjusted by false-discovery rate (FDR) correction. The hash (#) symbol is used to indicate comparisons with a WT control cohort (GCaMP6s; *nacre*) not derived from the experimental cross. * or #, adj P<0.05. ** or ##, adj P<0.01. *** or ###, adj P<0.001. **** or ####, adj P<0.0001.

First, at low-concentration PTZ, we saw a clear line-specific reduction in PTZ sensitivity affecting all *scn1lab^s^*^552^ conspecifics (**Fig 2B.i**), which is not observed in F0 crispant fish. For example, with respect to rate of classified seizure-like events, control animals from the *s552* line are dramatically reduced (wild-type: median 0/min, IQR 0-0.867; HET, 0.03/min, IQR 0-0.567) versus heterospecific wild-type controls (WT1: median 0.483/min, IQR 0.23-0.858, adj P = 0.0001 (versus WT), adj P <0.0001 (versus HET)). Control animals from crispant experiments did not differ in their sensitivity to PTZ (uninjected, median 0.767/min, IQR 0.467-2.03; NO CUT, 0.633/min, IQR 0.467-1.73) relative to heterospecific controls (adj P = 0.2 (versus uninjected), adj P =0.087 (versus NO CUT)). Given the partially shared genetic background between these lines, the reduction in sensitivity in the *scn1lab s552* conspecifics could be mediated either by a dominant effect of the previous genetic background (derived from the imported TL line) and/or parental effects of *scn1lab* LoF in mutant gametes.

Second, we see strong evidence of an *scn1lab*-related enhancement in PTZ induced seizures in both F0 crispant and stable *s552* fish (**Fig 2B.i**). For example, LOW/HIGH F0 crispant animals had elevated event rates (LOW, median 3.2/min, IQR 2.05-3.95; HIGH, 2.97/min, IQR 1.55-4.05) versus conspecifics (LOW versus uninjected, adj P = 0.01; HIGH versus uninjected, adj P = 0.03). Importantly, the enhanced sensitivity to PTZ in F0 crispant animals is even higher than observed in heterospecific wild-type controls (LOW versus WT1, adj P<0.0001; HIGH versus WT1, adj P = 0.0027). In contrast, *s552* HOM fish showed elevated event rates (median 0.233/min, IQR 0-2.9) relative to conspecifics (HOM versus WT, adj P = 0.0077; HOM versus HET, adj P > 0.0007), but were not different from heterospecific controls (HOM versus WT1, adj P =0.07). This again corroborates that the mechanism of the suppressive effect observed in the *s552* line is independent of larval genotype, and appears to attenuate the *scn1lab*-related phenotype in HOM animals. It is also worth mentioning that in F0 crispants, the *scn1lab*-related enhancement is detectable in fish with either LOW (0-50%) or HIGH (>50%) cutting, whereas only *s552* HOM (not HET) larvae showed this phenotype, suggesting that high cutting is not necessary to recapitulate a phenomenon that in stable lines appears to require biallelic disruption^16^.

Third, we next looked at changes in max velocity in response to low-concentration PTZ (**Fig 2B.ii**). In general, max velocity appears to be less informative than event rate, with no differences between conspecifics observed in either F0 crispant or stable *s552* larvae. Of note, despite showing a suppression of velocity and distance during spontaneous activity, the *s552* line shows no evidence for impairment in max velocity after exposure to PTZ, with all groups showing max velocities greater than 30 mm/s, similar to heterospecific controls. By contrast, the max velocity observed amongst F0 crispants was slightly reduced (uninjected, median 23.5 mm/s, IQR 20-32.6; LOW, 24.6 mm/s, IQR 19.2-27.6) versus heterospecific controls (uninjected versus WT1, adj P = 0.02, LOW versus WT1, adj P < 0.0001). Based on our experience with PTZ related seizures, this suggests that the seizure-like activity experienced by F0 crispants at low-concentration PTZ is in some ways similar to seizures seen in heterospecific controls at higher concentration PTZ (**cf. Fig 2C.ii**). The results of normalized calcium fluorescence per event (max deltaF/F0; **Fig 2B.iii**) are similar, with reductions in F0 crispants relative to *s552* and heterospecific controls and additional reductions in LOW and HIGH F0 crispants relative to conspecifics, which may reflect the higher rate of events in these larvae more typically observed at higher concentration PTZ (**cf. Fig 2C.iii**).

We next asked whether *scn1lab* LoF would alter the response to high-concentration PTZ (15mM) (**Fig 2C**). First, regarding the rate of classified seizure-like events, *s552* larvae showed the expected increase in the rate of events versus low-concentration PTZ (WT_PTZ2.5 vs. WT_PTZ15, adj P = 0.002; HET_PTZ2.5 vs. HET_PTZ15, adj P<0.0001; HOM_PTZ2.5 vs. HOM_PTZ15, adj P =0.009), but all conspecifics were still suppressed relative to heterospecific controls. In addition, the *scn1lab* related sensitivity in *s552* HOM larvae is no longer significantly different at high concentration PTZ (WT, median 0.55/min, IQR 0.033-2.26; HET, 0.4/min, IQR 0.033-2.27; HOM, 1.73/min, IQR 0.175-2.46)).

Second, in F0 crispants, conspecific controls showed the expected increase in seizure-like activity versus low-concentration PTZ (Uninjected_PTZ2.5 vs. Uninjected_PTZ15, adj P = 0.005; NO CUT_PTZ2.5 vs. NO CUT_PTZ15, adj P = 0.006), similar to heterospecific controls, but LOW and HIGH F0 crispants did not show higher rates (LOW _PTZ2.5 vs. LOW _PTZ15, adj P = 0.1889; HIGH_PTZ2.5 vs. HIGH_PTZ15, adj P = 0.185). In the case of HIGH F0 crispants, the event rates appear reduced (median 2.43/min, IQR 1.82-2.5) relative to conspecifics (uninjected: 3.067/min, IQR 2.57-3.37, adj P= 0.04; NO CUT: 3.1/min, IQR 2.13-3.7, adj P = 0.029). These observations again suggest that F0 crispant fish achieve more severe and frequent seizure-like activity at low-concentration PTZ, such that the effect is already saturated at higher PTZ concentrations, likely contributing to early lethality in these animals.

Third, both F0 crispant and stable *s552* lines showed *scn1lab*-related elevations in max velocity (Fig 2C.ii) and normalized calcium fluorescence (Fig 2C.iii) relative to conspecifics, which may be a mark of enhanced severity of seizures relative to conspecifics, despite similar event rates at this concentration.

In summary, we demonstrate that enhanced sensitivity to low-concentration PTZ is a robust correlate of *scn1lab* loss-of-function in F0 crispant and stable *s552* line, and show evidence for a still unexplained suppressive effect of the *s552* background on this phenotype.

### 3.3. Bootstrap simulations provide benchmarks for detecting scn1lab-related enhanced sensitivity to low-concentration PTZ in scn1lab F0 and scn1lab^s552^ in hypopigmented transgenic GCaMP6s zebrafish

Given our observations that neither the well-established *scn1lab^s^*^552^ line nor a separate *scn1lab* F0 crispant line show spontaneous seizure-like activity, further screens for spontaneous seizure-like activity under these conditions should proceed only with great caution. However, given the robust nature of the enhancement to low-concentration PTZ and its correspondence with *scn1lab* LoF, we foresee that reverse or forward genetic screens to detect gene-specific enhancements to low-concentration PTZ could be employed using the combined movement and fluorescence approach. To identify the optimal parameters for such screens, we performed two sets of bootstrap resampling simulations using the acquired datasets from *scn1lab* F0 and *scn1lab^s^*^552^ zebrafish. Using the robust strictly standardized mean difference (RSSMD) as a measure of effect size and variability, these calculations (3000 iterations, with replacement) compute the RSSMD threshold for detecting the observed enhancement in the rate of seizure-like events in a target group (*scn1lab* loss of function) versus a background group, as a function of bootstrap sample size (n=8-48) while maintaining 5% false positive rate (FPR). For F0 crispant simulations, we pooled all injected animals into the target group (i.e. no stratification) to mirror the real-life circumstances of an F0 screen where no filtering of the results based on the measured level of locus-specific cutting efficiency would be expected; uninjected conspecifics were defined as background. For reference, we also performed simulations with the *s552* data, with HOM animals defined as the target group, while WT and HET animals were pooled to form the background group. The results are shown as dual-axis plots (**Fig 3**) with RSSMD thresholds read as closed circles on the left axis, with associated true positive rates (TPR) read as open squares on the right axis.

**Figure 3.**
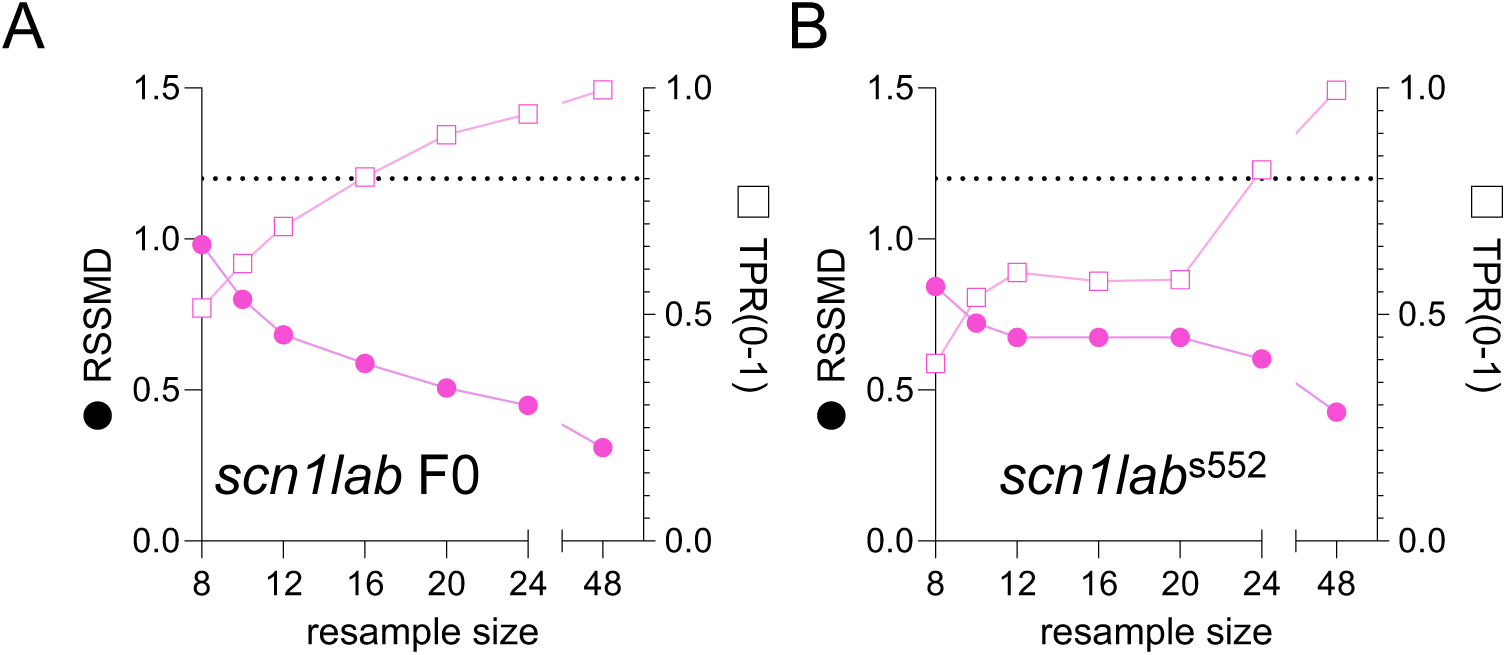
Bootstrap simulations provide benchmarks for detecting *scn1lab*-related enhanced sensitivity to low-concentration PTZ in *scn1lab* F0 and *scn1lab^s^*^552^ hypopigmented transgenic GCaMP6s zebrafish. **(A-B)** Bootstrap resampling simulations (3000 iterations, with replacement) from low-concentration PTZ for *scn1lab* F0 (A) and *scn1lab^s^*^552^ (B) zebrafish. For each bootstrap sample size N (x-axis), closed circles (left y-axis) are the robust strictly standardized mean difference (RSSMD) threshold required to limit the false positive rate (FPR) to 5% and open boxes (right y-axis) are the associated true positive rate (TPR) for detecting the *scn1lab*-related increase in PTZ sensitivity. Dashed reference lines indicate 80% TPR.

We observed that the enhanced PTZ sensitivity by the classified seizure-like event rate is detectable in F0 crispants (**Fig 3A**) with lower replicates (e.g. n=16, TPR 80%) versus *scn1lab^s^*^552^ (e.g. n=24 required to achieve TPR 80%; **Fig 3B**). This is interesting considering the statistical magnitude of effect for these differences (see **Fig 2Bi**), but can be explained by the fact that in F0, the control groups (uninjected and NO CUT) each had 13-15 animals, which is below the minimum sample size suggested by the analysis (N=16, at TPR 80%). In *s552* larvae, the controls (WT/HET) had 31-55 animals – well above the minimum sample size suggested by the analysis (N=24 and closer to N=48, at which 100% TPR is achieved).

Ultimately, both F0 crispants and stable lines appear suitable for screening at higher sample sizes using combined movement and fluorescent profiling, with crispants demonstrating a slight advantage with respect to the minimum number of replicates necessary.

## 4. Discussion

Advanced analysis of brain activity from model organisms in states of health and disease benefits from the combination of different stable lines, including those expressing disease-relevant mutations and/or transgenic lines expressing genetically encoded calcium indicators (such as GCaMP6s) among other tools. However, the effects of these combinations or changes in genetic background on the phenotype under investigation are not always rigorously assessed or controlled in zebrafish. This is a critical issue for the rigor and reproducibility of animal studies, which undergirds the legitimacy by which the pathophysiological mechanisms associated with disease alleles may be dissected through the use of tools generated on different genetic backgrounds. Although these issues are well-known in the rodent literature ^23^, comparatively little attention has been paid in the zebrafish literature^8^.

In the present study, we attempted to evaluate spontaneous seizure-like activity and proconvulsant-related seizure-like activity associated with loss-of-function in the well-characterized *scn1lab* gene in zebrafish expressing a genetically encoded calcium indicator (elavl3:GCaMP6s) with *nacre* hypopigmentation phenotype using combined movement and fluorescence profiling (summarized in **Table 1**), but encountered several challenges which highlight the importance of understanding the implications of seemingly routine genetic manipulations on the phenotype under study.

**Table 1.** Summary of major findings

|  | *scn1lab* F0 | *scn1lab*<sup>s552</sup> |
| --- | --- | --- |
| **Background** | AB | 50:50 AB:TL |
| **Rapid swimming phenotype** | Not observed | Not observed |
| **Spontaneous max velocity** | ↓ vs conspecifics<br>↓↓ vs unrelated WT | ↑ vs conspecifics<br>↓↓ vs unrelated WT |
| **Normalized calcium fluorescence per bout**<br>(average max dF/F0) | ↓ vs conspecifics<br>↓/no Δ vs unrelated WT | ↑ vs conspecifics<br>↓/no Δ vs unrelated WT |
| **Spontaneous bout rate** (rate of unclassified calcium events at baseline) | ↓ vs conspecifics<br>No Δ vs unrelated WT | ↑ vs conspecifics<br>No Δ vs unrelated WT |
| **Sensitivity to low-dose PTZ**<br>(rate of classified seizure events in response to PTZ 2.5mM) | <u>↑ vs conspecifics</u><br>↑↑ vs unrelated WT<br>No Δ (uninjected) vs unrelated WT | <u>↑ vs conspecifics</u><br>No Δ vs unrelated WT<br>↓↓ (WT/HET) vs unrelated WT |

### 4.1. No detectable spontaneous seizure-like phenotype in scn1lab lines based on combined movement and fluorescence measurements

By conducting our experiments using a GCaMP6s;*nacre* line, we accidentally discovered conditions that suppress the rapid swimming phenotype associated with the well-known epilepsy allele *scn1lab^s^*^552^, in addition to the effects of *scn1lab* LoF in F0 crispant fish, representing two independent lines of evidence. Although we do not directly model the *s552* missense variant (M1208R) in *scn1lab* F0 crispants, the nature of the disruption in F0 crispants would be expected to be stronger than that of *s552*, and yet it also does not result in spontaneous seizure-like activity. This is surprising because the phenotype associated with *scn1lab^s^*^552^ has been so well-established^5,16^.

Indeed, there are many reports in the literature affirming the locomotor phenotype of *s552*^24^, or in which alternative stable *scn1lab* LoF alleles^17,18^, morpholino knock-downs^24^ or F0 crispants^25^ are generated and shown also to display similar locomotor phenotypes. The *s552* line maintained in Baraban’s group is on TL^5^, as is ours, whereas the Tiraboschi group implies that its novel *scn1lab* KO allele is on AB^17^; other authors did not report the genetic background used. No authors use the GCaMP6;*nacre* line utilized in this study, which is maintained on AB. All lines shown to have rapid swimming or increased locomotor activity also showed other abnormalities by tectal LFP and/or calcium fluorescence. There were no reports of rapid swimming phenotype being lost after combination with other lines, though it is not clear if this was assessed in the two studies using calcium fluorescence^20,24^, and it is interesting to note that the Tiraboschi group did not report rapid swimming but rather increased distance traveled in *scn1lab* KO animals on AB background. In summary, there is incomplete information from the literature to determine the extent to which genetic background or the combination of other zebrafish transgenic or mutant lines has contributed to the rapid swimming phenotype reported by other authors in association with *scn1lab* LoF mutations.

We speculate that our findings could be related to an effect of one or more factors unique to our study, including genetic background, *nacre* hypopigmentation, or the GCaMP6s transgenic line. First, a dominant suppressive effect of the AB background on which the GCaMP6s; *nacre* line is maintained could account for the effect in both stable *s552* and F0 crispant fish. Crispant and heterospecific GCaMP6s; *nacre* control larvae were derived directly from this AB-derived line, while F2 *scn1lab^s^*^552^ larvae used for experiments are expected to be 50:50 AB:TL, suggesting one or more genetic modifiers from AB may act to suppress *scn1lab*-related hyperexcitability.

Second, it is possible that the *nacre* pigmentation defect, or the genetic processes leading to it, could play a role. It is hard to ignore that *scn1lab* HOM have a well-known but poorly understood hyperpigmentation phenotype (seen in multiple lines, including *s552*^16,19,24^ and those of others^17,24^), but it has never been asserted to have a causal role in *scn1lab-*related hyperexcitability. Naturally, neither *s552* or F0 crispant animals in our study show hyperpigmentation since the *nacre* phenotype results from an absence of melanophores due to recessive loss-of-function in the *mitfa* (aka *nacre*) gene^4^, suggesting that ablation of melanophores may be a candidate mechanism for suppression of *scn1lab*-related spontaneous seizure-like activity. Since the *mitfa* locus is commonly combined with other loci to generate more extensive pigmentation deficits such as *casper* (*mitfa^w^*^2^*^/w^*^2^*;roy^a^*^9^*^/a^*^9^) and *crystal* (*mitfa^w^*^2^*^/w^*^2^*;alb^b^*^4^*^/b^*^4^*;roy^a^*^9^*^/a^*^9^), any undesired effects of *nacre* on the *scn1lab* phenotype may be highly relevant to other hypopigmentation combinations as well. To the best of our knowledge, no other studies involving *scn1lab* utilize pigmentation mutants. The use of 1-phenyl 2-thiourea (PTU)--a chemical inhibitor of tyrosinase, which inhibits melanin production but preserves melanophores^26^ -- has been reported twice^20,24^ though its effect on rapid swimming was not assessed. The use of PTU is generally regarded as having greater deleterious neurological effects^26–28^ compared to pigmentation mutants. Meanwhile, F0 *scn1lab* crispants that harbor concomitant acute KO in the *tyr* gene still have rapid swimming^25^, suggesting that lack of functional tyrosinase enzyme is not sufficient to suppress.

Third, we consider it is less likely to be related to GCaMP itself. In the mouse literature, a consistent pro-epileptic phenotype has been reported with specific GCaMP6s transgenic lines^23^, due to what the authors argue is an effect of widespread GCaMP6 expression specifically during brain development, as opposed to the genetic background or toxicity from Cre or tTA used in these lines. Perhaps if GCaMP6s can have pro-epileptic effects in one context, compensatory anti-epileptic effects may be triggered during zebrafish development, but this may be the least compelling explanation for our findings. In the context of *scn1lab* zebrafish, there are two other examples of *scn1lab* mutants with seizure-like activity of some kind in combination with elavl3:GCaMP5, suggesting that GECIs do not categorically suppress *scn1lab*-related seizure activity^20,24^. However, there are no other relevant studies reported using the transgenic GCaMP6s line (employed here), so a line-specific phenotype due to insertion effects of the transgene^29,30^ or linked modifier loci cannot yet be excluded.

Future experiments should explore the molecular mechanism of this suppression, to test its dependency on specific lines used here and to test specifically whether hypopigmentation suppresses *scn1lab* pathophysiology. These studies would shed more light on the factors contributing to variability in preclinical zebrafish models of epilepsy, and may identify genetic modifiers with clinical relevance to Dravet syndrome.

### 4.2. Enhanced sensitivity to PTZ in scn1lab lines

At the same time, we also see recapitulation of enhanced susceptibility to PTZ in a manner dependent on *scn1lab* LoF. This is evidenced by elevated rate of PTZ-like seizure activity after exposure to acute low-concentration PTZ (2.5 mM) in both *s552*-HOM and F0 crispant fish, as quantified using the combined movement and fluorescence profiling and the PTZ M+F classifier. These findings suggest that although *scn1lab* deficiency does not result in spontaneous PTZ-like seizures under the conditions reported, nevertheless *scn1lab* deficiency alters excitatory-inhibitory balance in a manner that lowers the seizure threshold provoked by GABA_A_R antagonism, perhaps due to impaired inhibitory versus enhanced excitatory synaptic transmission. This finding is consistent with other reports of enhanced sensitivity to PTZ in a model of *scn1lab*^18^, in addition to evidence of reduced whole organism GABA levels in *scn1lab* zebrafish^18^. Meanwhile, we show remarkably low percentage of measured cutting (0-50%) in *scn1lab* was sufficient to generate a prominent PTZ phenotype in F0 crispants, justifying the use of F0 crispants in reverse genetic screening for seizure-related phenotypes.

### 4.3. Could scn1lab lines still have spontaneous seizures that are less severe?

Taking both findings (4.1 and 4.2) together, we concede that it is possible that *scn1lab* LoF mutants in this study may yet have evidence of spontaneous seizures – perhaps less severe, and with minimal movement -- by other modalities not assessed here, such as tectal LFP. We expected to be able to detect milder seizures using movement or calcium fluorescence by way of a HOM-specific classifier, and we did demonstrate type 0 events detected by this classifier occur at elevated rates compared to conspecific controls. However, these events are not easily distinguished by movement and fluorescence criteria from physiological events, at least under the conditions reported here, limiting their utility as a read-out on this platform. Of note, we can confidently exclude the possibility that suppression of rapid swimming occurs due to a deficit in movement generation, as both *s552* and F0 crispants are capable of rapid movements (>20mm/sec) in response to PTZ. Future investigations should determine whether genetic mutants on different background lines may have seizure-like activity that is more amenable to detection on this platform.

### 4.4. Genetic suppression of PTZ sensitivity in larvae derived from scn1lab^s552/+^ matings

Last, we also report evidence for a second mode of suppression in the *scn1lab*-*s552* line. This phenomenon is expressed as a reduction in the rate of seizure-like calcium events induced by low-concentration PTZ (2.5mM) and high-concentration PTZ (15mM) across conspecific animals generated from *scn1lab^s^*^552/+^ parents. We believe this is distinct from the mechanism suppressing the rapid swimming phenotype in *s552* and F0 crispants, since conspecific control larvae from F0 crispant experiments and heterospecific controls did not show reduced PTZ sensitivity.

The mechanism is also unclear. A dominant effect of the 50% TL genetic background remaining from the imported *s552* line (see **Section 4.1**) could explain why the phenotype is only observed in larvae derived from *scn1lab^s^*^552^ parents. A transcriptional adaptation (TA) response^12^ (see **Introduction**) is unlikely as *s552* is a missense variant and not anticipated to induce NMD. Another possibility is a maternal effect due to *scn1lab*-related misregulation of paralogous sodium channel genes in the maternal zygote. Sodium channel expression and function are well-known to be subject to homeostatic regulation during development^31^. Based on publicly available mRNA expression data from zebrafish development^32^, transcripts from several voltage-gated sodium genes are present at the earliest zygotic time-points (including *scn1bb*, *scn1laa* and others, but not *scn1lab* itself; **Supplemental Fig 3**) and might be candidate genes whose putative misregulation in the setting of maternal *scn1lab* LoF alters larval sensitivity to PTZ. In support of this possibility, an experimental over-expression of the paralogous *scn1laa* only during the first 24hrs of development was sufficient to cause an epileptiform phenotype at later time-points and to worsen the phenotype of *scn1lab* LoF larvae^33^, demonstrating that early regulation of voltage-gated sodium channel genes is highly influential on later seizure-related phenotypes. Future work should explore the molecular mechanism of this suppression and whether it relates to misregulation of sodium channels in the context of *scn1lab* LoF.

To the best of our knowledge, this is the first report of a suppressive phenomenon related to the *scn1lab^s^*^552^ allele affecting WT offspring. One reason it has not been previously reported may be because the use of both conspecific and heterospecific controls is not routine. It is also an anti-epileptic phenotype, which requires proconvulsant exposure to detect in WT offspring, due to the lack of spontaneous seizure-like activity. Nevertheless, data from Griffin et al^34^ suggest that the prevalence of parental (likely maternal) effects in association with putative epilepsy genes in zebrafish may be greater than has been formally recognized. In this paper, the authors generated 40 lines via CRISPR/Cas9 corresponding to homologs of human epilepsy genes, and reported a low prevalence of spontaneous seizure-like activity among HOM animals (compared to conspecifics), but widely divergent findings between the WT conspecifics of different presumably congenic stable lines. Some WT animals showed a greater amount of epileptiform abnormality from tectal electrophysiological recordings than conspecific homozygotes or heterospecific WTs (for example, *scn1ba*, *scn8aa,* and several others^34^). Limited explanation for the WT phenotypes is offered by the authors, but an effect of GC may be a compelling explanation that should be assessed in future endeavors to model genetic epilepsy in zebrafish using stable alleles.

## 5. Data and code availability statement

All of the data generated in the present study and MATLAB/R code are available upon request.

## 6. Acknowledgements

The authors would like to thank Lee Barrett for technical assistance with the FDSS7000EX, Guoqi Zhang for assistance in zebrafish husbandry, and members of the Poduri lab including Christopher LaCoursiere for helpful discussions.

## 7. Author contributions using the CRediT taxonomy

CM: Conceptualization, Data curation, Formal analysis, Funding acquisition, Investigation, Methodology, Software, Visualization, Supervision, Writing – original draft, Writing – review and editing.

CB: Investigation, Writing – review.

AP: Funding acquisition, Supervision, Writing – review and editing.

## 11. Supplementary Material

### 11.1. Supplementary Methods

#### 10.1.1. Supervised machine learning for event classification in *scn1lab^s^*^552^ HOM fish

To evaluate whether calcium events from *scn1lab^s^*^552^ HOM fish might have unique features that distinguish them from events occurring in WT fish, a logistic classifier was trained using the same approach as in **Section 2.7**. using events from all *s552* HOM animals versus *s552* WT controls, 70:30 train:test split, and 5-fold cross-validation. The model formula for the classifier (referred to as the “scn1lab M+F” model) was identical to that of the previously described “PTZ M+F” classifier^6^. Specifically, the model formula used was: Conditions_names ∼ (MaxIntensity_F_centroid + MaxIntensity_F_F0_centroid + distance_xy_mm + duration_sec) * (maxRange_2 + totalCentroidSize_mode_mm2). Data were divided into 70:30 train:test split, with 5-fold cross validation, alpha range: 0,0.5, 1, lambda range: 0.1, 1, 10, and metric = “accuracy”. Model performance was evaluated using package MLeval.

### 11.2. Supplemental Figures

**Supplementary Figure 1.**
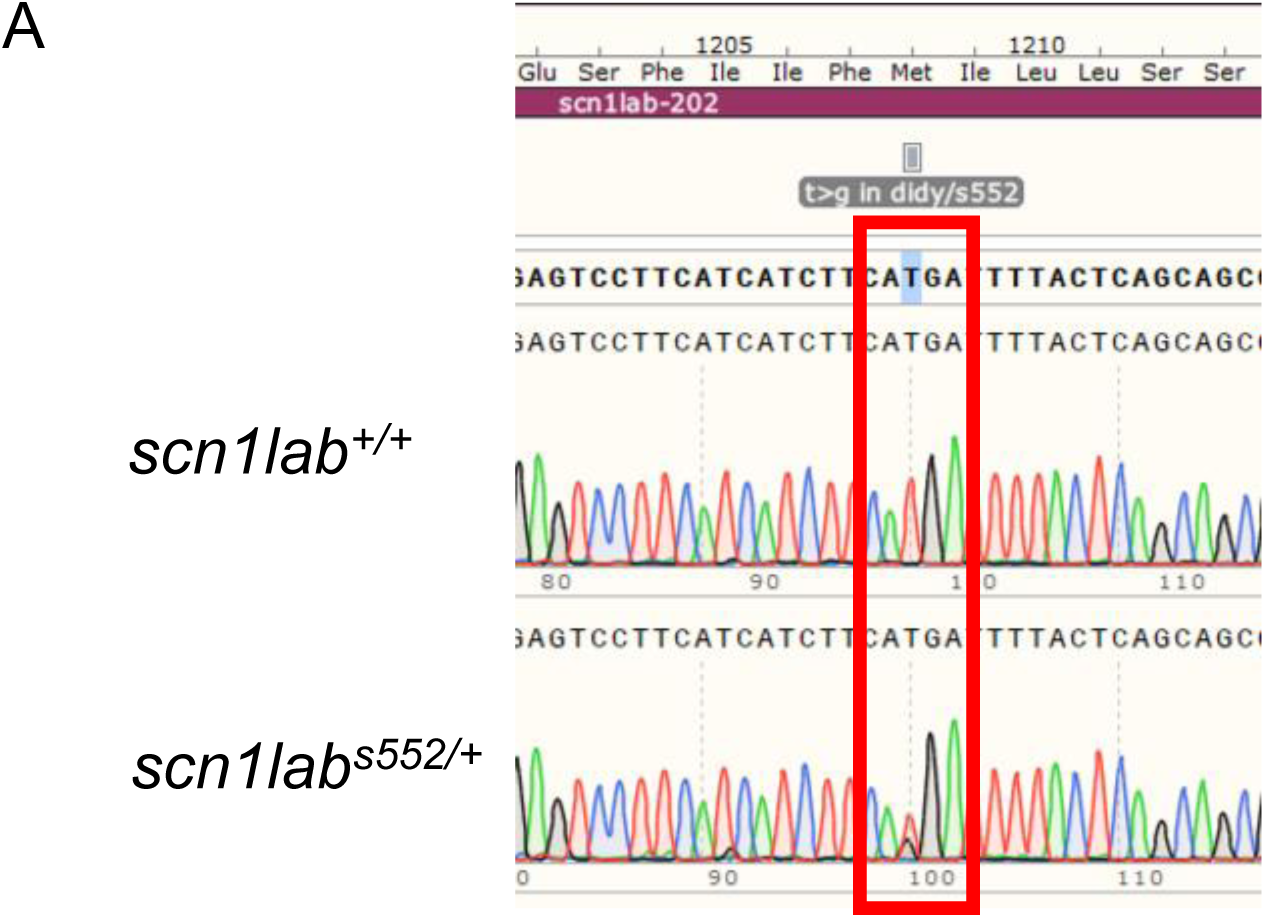
Generation of *scn1lab^s^*^552^ larvae. (A) Representative example of Sanger sequencing from PCR genotyping of *scn1lab^s^*^552^ larvae, demonstrating the expected c.T>G variant in a heterozygote larvae (lower) vs wildtype (middle) compared to reference sequence (upper)

**Supplementary Figure 2.**
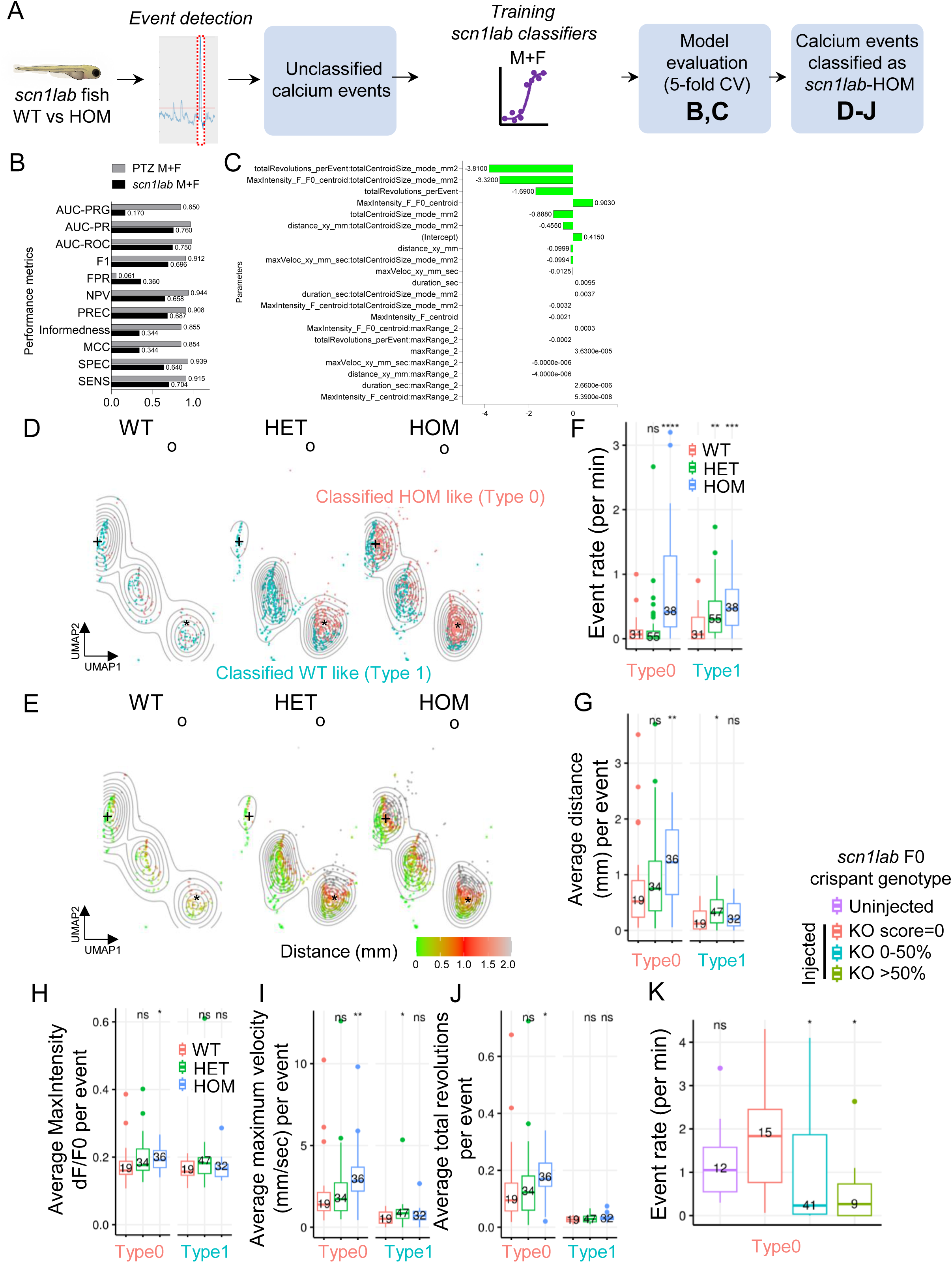
Elastic net logistic classifier trained on *scn1lab^s^*^552^ HOM larvae weakly detects an event type enriched in s552 HOM larvae. **(A)** Overview of approach and figure. **(B)** Comparison of performance metrics from *scn1lab* M+F classifier versus the previously published PTZ M+F classifier trained on PTZ-induced seizure activity. (**C**) Parameter tuning from *scn1lab* M+F classifier. **(D-E)** Uniform manifold approximation and projection (UMAP) representation of pooled individual events from *scn1lab^s^*^552^ conspecifics, color coded by classification (D) or average distance (E). **(F-G)** Group-wise quantification of event rate (F) and average distance (G) from larvae in (D-E), stratified by classified event type (Type 0 vs Type 1). **(H-J)** Group-wise quantification of normalized calcium fluorescence (H), max velocity (I), and total revolutions (J) for *scn1lab^s^*^552^ larvae, stratified by classified event type. (**K**) Event rates derived from events classified as Type 0 by the *scn1lab* M+F classifier in *scn1lab* F0 crispant fish. Data are group-wise Tukey box-plots of subject-level averages of all subject-specific events. Reported p-values are derived from Wilcoxon rank sum test, and adjusted by false-discovery rate (FDR) correction. *, adj P<0.05. **, adj P<0.01. ***, adj P<0.001. ****, adj P<0.0001.

**Supplementary Figure 3.**
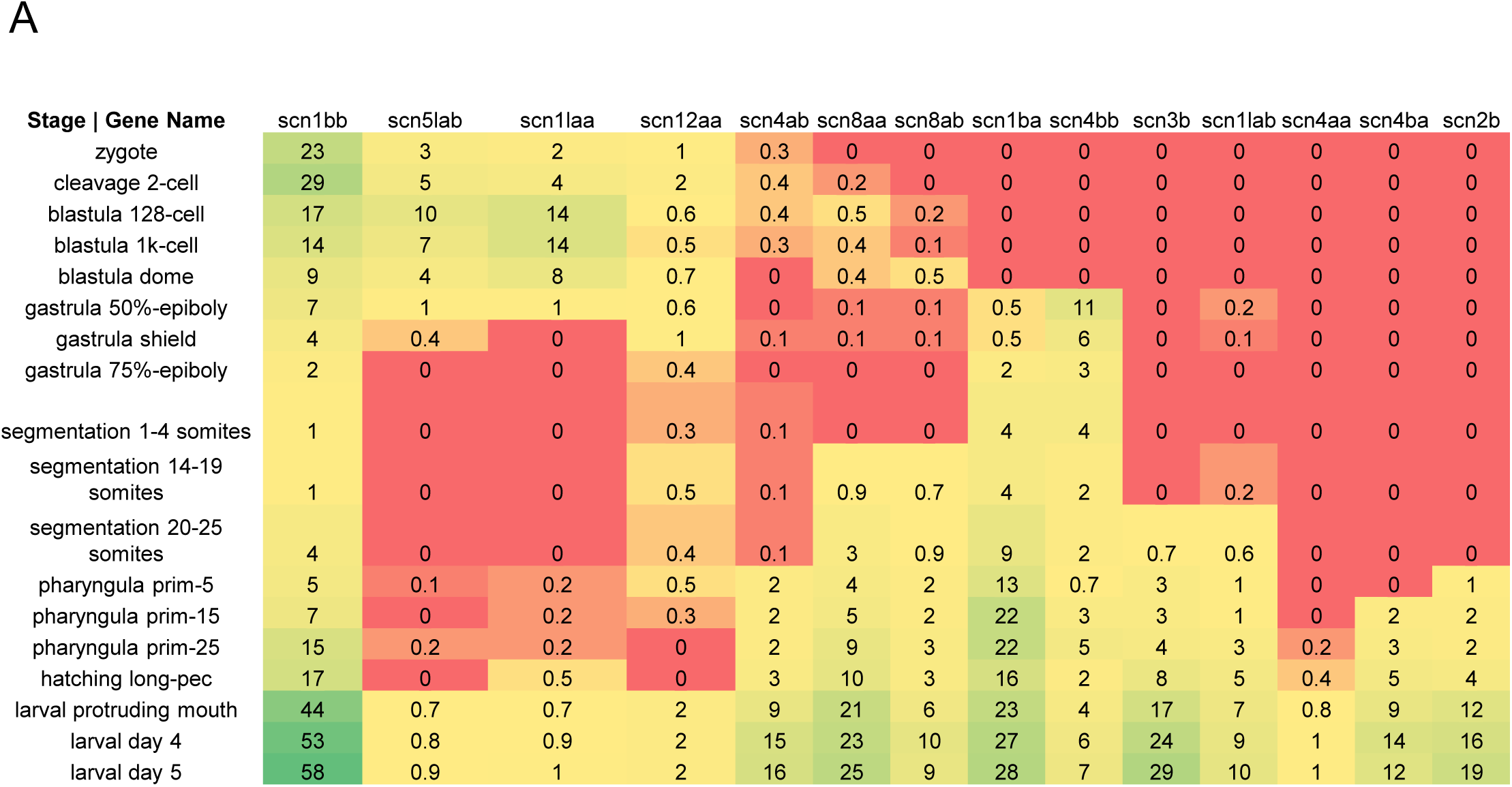
Expression of sodium channels genes across larval zebrafish developmental stages. Data are transcripts per million (TPM) derived from mRNA-seq, as reported by White RJ, Collins JE, Sealy IM, Wali N, Dooley CM et al. (2017) A high-resolution mRNA expression time course of embryonic development in zebrafish. Colors are scaled continuously from minimum (red), median (yellow), to max (green).

## References

1. Gawel, K., Langlois, M., Martins, T., van der Ent, W., Tiraboschi, E., Jacmin, M., Crawford, A.D., and Esguerra, C.V. (2020). Seizing the moment: Zebrafish epilepsy models. Neurosci. Biobehav. Rev. 116, 1–20. 10.1016/j.neubiorev.2020.06.010.

2. LaCoursiere, C.M., Ullmann, J.F.P., Koh, H.Y., Turner, L., Baker, C.M., Robens, B., Shao, W., Rotenberg, A., McGraw, C.M., and Poduri, A. (2024). Zebrafish models of candidate human epilepsy-associated genes provide evidence of hyperexcitability. Preprint at bioRxiv, 10.1101/2024.02.07.579190.

3. Vladimirov, N., Mu, Y., Kawashima, T., Bennett, D.V., Yang, C.-T., Looger, L.L., Keller, P.J., Freeman, J., and Ahrens, M.B. (2014). Light-sheet functional imaging in fictively behaving zebrafish. Nat. Methods 11, 883–884. 10.1038/nmeth.3040.

4. Lister, J.A., Robertson, C.P., Lepage, T., Johnson, S.L., and Raible, D.W. (1999). nacre encodes a zebrafish microphthalmia-related protein that regulates neural-crest-derived pigment cell fate. Development 126, 3757–3767. 10.1242/dev.126.17.3757.

5. Dinday, M.T., and Baraban, S.C. (2015). Large-Scale Phenotype-Based Antiepileptic Drug Screening in a Zebrafish Model of Dravet Syndrome. eNeuro 2. 10.1523/ENEURO.0068-15.2015.

6. McGraw, C.M., and Poduri, A. (2024). Machine learning enables high-throughput, low-replicate screening for novel anti-seizure targets and compounds using combined movement and calcium fluorescence in larval zebrafish. Preprint at bioRxiv, 10.1101/2024.08.01.606228.

7. Trevarrow, B., and Robison, B. (2004). Genetic Backgrounds, Standard Lines, and Husbandry of Zebrafish. In Methods in Cell Biology The Zebrafish: Genetics, Genomics, and Informatics. (Academic Press), pp. 599–616. 10.1016/S0091-679X(04)77032-6.

8. Crim, M.J., and Lawrence, C. (2021). A fish is not a mouse: understanding differences in background genetics is critical for reproducibility. Lab Anim. 50, 19–25. 10.1038/s41684-020-00683-x.

9. Löscher, W., Ferland, R.J., and Ferraro, T.N. (2017). The relevance of inter- and intrastrain differences in mice and rats and their implications for models of seizures and epilepsy. Epilepsy Behav. 73, 214–235. 10.1016/j.yebeh.2017.05.040.

10. El-Brolosy, M.A., and Stainier, D.Y.R. (2017). Genetic compensation: A phenomenon in search of mechanisms. PLOS Genet. 13, e1006780. 10.1371/journal.pgen.1006780.

11. El-Brolosy, M.A., Kontarakis, Z., Rossi, A., Kuenne, C., Günther, S., Fukuda, N., Kikhi, K., Boezio, G.L.M., Takacs, C., Lai, S.-L., et al. (2019). Genetic compensation triggered by mutant mRNA degradation. Nature 568, 193–197. 10.1038/s41586-019-1064-z.

12. Jiang, Z., El-Brolosy, M.A., Serobyan, V., Welker, J.M., Retzer, N., Dooley, C.M., Jakutis, G., Juan, T., Fukuda, N., Maischein, H.-M., et al. (2022). Parental mutations influence wild-type offspring via transcriptional adaptation. Sci. Adv. 8, eabj2029. 10.1126/sciadv.abj2029.

13. Pelegri, F. (2003). Maternal factors in zebrafish development. Dev. Dyn. 228, 535–554. 10.1002/dvdy.10390.

14. Buglo, E., Sarmiento, E., Martuscelli, N.B., Sant, D.W., Danzi, M.C., Abrams, A.J., Dallman, J.E., and Züchner, S. (2020). Genetic compensation in a stable slc25a46 mutant zebrafish: A case for using F0 CRISPR mutagenesis to study phenotypes caused by inherited disease. PLOS ONE 15, e0230566. 10.1371/journal.pone.0230566.

15. Dravet, C. (2011). The core Dravet syndrome phenotype. Epilepsia 52, 3–9. 10.1111/j.1528-1167.2011.02994.x.

16. Baraban, S.C., Dinday, M.T., and Hortopan, G.A. (2013). Drug screening in Scn1a zebrafish mutant identifies clemizole as a potential Dravet syndrome treatment. Nat. Commun. 4, 2410. 10.1038/ncomms3410.

17. Tiraboschi, E., Martina, S., van der Ent, W., Grzyb, K., Gawel, K., Cordero-Maldonado, M.L., Poovathingal, S.K., Heintz, S., Satheesh, S.V., Brattespe, J., et al. (2020). New insights into the early mechanisms of epileptogenesis in a zebrafish model of Dravet syndrome. Epilepsia 61, 549–560. 10.1111/epi.16456.

18. Weuring, W.J., Singh, S., Volkers, L., Rook, M.B., Slot, R.H. van ‘t, Bosma, M., Inserra, M., Vetter, I., Verhoeven-Duif, N.M., Braun, K.P.J., et al. (2020). NaV1.1 and NaV1.6 selective compounds reduce the behavior phenotype and epileptiform activity in a novel zebrafish model for Dravet Syndrome. PLOS ONE 15, e0219106. 10.1371/journal.pone.0219106.

19. Schoonheim, P.J., Arrenberg, A.B., Bene, F.D., and Baier, H. (2010). Optogenetic Localization and Genetic Perturbation of Saccade-Generating Neurons in Zebrafish. J. Neurosci. 30, 7111–7120. 10.1523/JNEUROSCI.5193-09.2010.

20. Ghannad-Rezaie, M., Eimon, P.M., Wu, Y., and Yanik, M.F. (2019). Engineering brain activity patterns by neuromodulator polytherapy for treatment of disorders. Nat. Commun. 10, 2620. 10.1038/s41467-019-10541-1.

21. Labun, K., Montague, T.G., Krause, M., Torres Cleuren, Y.N., Tjeldnes, H., and Valen, E. (2019). CHOPCHOP v3: expanding the CRISPR web toolbox beyond genome editing. Nucleic Acids Res. 47, W171–W174. 10.1093/nar/gkz365.

22. Conant, D., Hsiau, T., Rossi, N., Oki, J., Maures, T., Waite, K., Yang, J., Joshi, S., Kelso, R., Holden, K., et al. (2022). Inference of CRISPR Edits from Sanger Trace Data. CRISPR J. 5, 123–130. 10.1089/crispr.2021.0113.

23. Steinmetz, N.A., Buetfering, C., Lecoq, J., Lee, C.R., Peters, A.J., Jacobs, E.A.K., Coen, P., Ollerenshaw, D.R., Valley, M.T., Vries, S.E.J. de, et al. (2017). Aberrant Cortical Activity in Multiple GCaMP6-Expressing Transgenic Mouse Lines. eNeuro 4. 10.1523/ENEURO.0207-17.2017.

24. Brenet, A., Hassan-Abdi, R., Somkhit, J., Yanicostas, C., and Soussi-Yanicostas, N. (2019). Defective Excitatory/Inhibitory Synaptic Balance and Increased Neuron Apoptosis in a Zebrafish Model of Dravet Syndrome. Cells 8, 1199. 10.3390/cells8101199.

25. Locubiche, S., Ordóñez, V., Abad, E., Scotto di Mase, M., Di Donato, V., and De Santis, F. (2024). A Zebrafish-Based Platform for High-Throughput Epilepsy Modeling and Drug Screening in F0. Int. J. Mol. Sci. 25, 2991. 10.3390/ijms25052991.

26. Li, Z., Ptak, D., Zhang, L., Walls, E.K., Zhong, W., and Leung, Y.F. (2012). Phenylthiourea Specifically Reduces Zebrafish Eye Size. PLOS ONE 7, e40132. 10.1371/journal.pone.0040132.

27. Antinucci, P., and Hindges, R. (2016). A crystal-clear zebrafish for in vivo imaging. Sci. Rep. 6, 29490. 10.1038/srep29490.

28. Chen, X.-K., Kwan, J.S.-K., Chang, R.C.-C., and Ma, A.C.-H. (2021). 1-phenyl 2-thiourea (PTU) activates autophagy in zebrafish embryos. Autophagy 17, 1222–1231. 10.1080/15548627.2020.1755119.

29. Goodwin, L.O., Splinter, E., Davis, T.L., Urban, R., He, H., Braun, R.E., Chesler, E.J., Kumar, V., Min, M. van, Ndukum, J., et al. (2019). Large-scale discovery of mouse transgenic integration sites reveals frequent structural variation and insertional mutagenesis. Genome Res. 29, 494–505. 10.1101/gr.233866.117.

30. Jacobsen, J.C., Erdin, S., Chiang, C., Hanscom, C., Handley, R.R., Barker, D.D., Stortchevoi, A., Blumenthal, I., Reid, S.J., Snell, R.G., et al. (2017). Potential molecular consequences of transgene integration: The R6/2 mouse example. Sci. Rep. 7, 41120. 10.1038/srep41120.

31. Schulz, D.J., Temporal, S., Barry, D.M., and Garcia, M.L. (2008). Mechanisms of voltage-gated ion channel regulation: from gene expression to localization. Cell. Mol. Life Sci. 65, 2215–2231. 10.1007/s00018-008-8060-z.

32. White, R.J., Collins, J.E., Sealy, I.M., Wali, N., Dooley, C.M., Digby, Z., Stemple, D.L., Murphy, D.N., Billis, K., Hourlier, T., et al. (2017). A high-resolution mRNA expression time course of embryonic development in zebrafish. eLife 6, e30860. 10.7554/elife.30860.

33. Weuring, W.J., Dilevska, I., Hoekman, J., van de Vondervoort, J., Koetsier, M., van ’t Slot, R.H., Braun, K.P.J., and Koeleman, B.P.C. (2021). CRISPRa-Mediated Upregulation of scn1laa During Early Development Causes Epileptiform Activity and dCas9-Associated Toxicity. CRISPR J. 4, 575–582. 10.1089/crispr.2021.0013.

34. Griffin, A., Carpenter, C., Liu, J., Paterno, R., Grone, B., Hamling, K., Moog, M., Dinday, M.T., Figueroa, F., Anvar, M., et al. (2021). Phenotypic analysis of catastrophic childhood epilepsy genes. Commun. Biol. 4, 1–13. 10.1038/s42003-021-02221-y.

